# Mathematical Modelling of Actin treadmill in *Apicomplexans*

**DOI:** 10.1101/258913

**Authors:** Mahendra Prajapat, Samridhi Pathak, Ricka Gauba, Avinash Kale, Supreet Saini

## Abstract

*Plasmodium* parasite, a representative member of phylum *Apicomplexa* is a causative agent of malaria in human as well as other animals. To infect host cells, *Plasmodium* first finds receptors on the host cell surface, then binds specifically, and finally penetrates host cell membrane to acquire the host cellular resources. The motility for moving on the cell surface is equipped by the precise and tight control of actin treadmill. Several regulators are required to achieve precision and robustness in the control of actin treadmill. However, the mechanistic detail of the treadmill regulatory network and the cross-talk among regulators are not well understood. We developed a stochastic model of treadmill regulation and explored the dynamics of filament growth, nucleation time, and elongation time. Our study mainly highlighted on how and what helps cells to maintain an average size of the actin filaments within a species. This is particularly important, since, excessive growth of filament can lead to cell lysis. Moreover, we also explore how the regulators interact to fine-tune the control elements in the actin treadmill.

## Introduction

Over the years, the process of building cytoskeletal filaments, also known as treadmill, has been studied in various contextssuch as molecular and cellular motility (1). In principle, the functioning of treadmill appears on continuously growing cytoskeletal filaments, the actin filaments and microtubules, in well controlled manner (1, 2). The single-cell eukaryotic organisms such as the members of *Apicomplexa* phylum use actin filaments dependent cellular motility to move in their surroundings (2, 3). A large number of parasitic protists from the *Apicomplexa* phylum are well known to cause a wide range of diseases in their respective hosts (1, 4, 5). The hallmark feature of these parasites is to acquire a complex structure called apical complex (5), which helps them penetrating the host cell membrane. *Plasmodium*, an obligate endoparasite, a causative agent of malaria in human and many other animals, is well known member of the phylum *Apicomplexa* (6). *Plasmodium* invades the host cells by receptor specific penetration (6), and thereafter, hijacks the host cell resources to further grow and reproduce its clone-mates. The search of appropriate receptors on the host cell surface equipped by the actin treadmill (6–9). Hence, the accuracy and speed of the *Plasmodium’s* motility has an important role in establishment of the early infection and is determined by the precise and robust control of actin treadmill.

Actin filaments grow in length at one end and shrink at the other end by continuous addition and removal of actin monomers respectively. It results in the filament apparently moving across the cytosol (10). The filament end where monomers are added is called as barbed (+) end and the other end where monomers are removed is called as pointed (-) end. In general, the growth of filaments occurs when the rate of addition at barbed end is comparatively higher than the rate of removal at pointed end (10). This difference in rates in equilibrium, which is constant in particular species, determines the size of the actin filament thus, in a single species, the average filament size is constant (10, 11). It has not been yet explored that how and what maintains the difference in the rates of polymerization and depolymerization in order to maintain average size of the filamentsconstant in a particular species.

A number of regulators, such as Formin (FH), Profilin, ADF (actin depolymerizing factor), CAP (cyclase associated protein), Capping protein, and Coronin protein (6, 12–14), control the treadmilling process and provide a sensible, timely correct, and efficient motility (6). The recent advancements in the field of structural biology, biophysics, biochemical and living cell imaging have significantly boosted our understanding of actin treadmill and its underlying molecular mechanism (15).However, the mechanistic details of the treadmill control and the interlay among the regulators are still not well understood.

To address these questions, we develop a mathematical model using *Plasmodium*as a model organism. In our model, we write ordinary differential equations for the biochemical interactions involved in the actin treadmill. The model was then simulated using Gillespie’s stochastic algorithm and the dynamics of treadmill was analyzed in various conditions. Our simulations highlight the elements of the regulatory network, which enable precise control of treadmill dynamics, and most importantly, limit the number of actin monomers that can get assembled in a single filament. This is particularly important, since, excessive growth of filament can lead to cell lysis. However, to avoid such a scenario, the control of the system is so tuned that the filament size cannot exceed an upper threshold, irrespective of regulator, or monomer concentration. Moreover, our model also predicts that capping protein helps preserving the actin dimers and trimers for the next consecutive treadmills even if the resources are limited.

## Method and model development

Using *Plasmodium* as a model organism, we developed a mathematical model of control mechanism of actin treadmill by writing ordinary differential equation for biochemical interactions involved in both nucleation and elongation (Equations 1–40). We simulated different variants of the model (as discussed below)using Gillespie’s algorithm to account for the stochasticity (16–18). The values of parameters in the model were chosen to fit the observed treadmill kinetics reported in the existing literature (7, 10, 19–25). All simulations were run for 1000 cells simultaneously and then the simulation dynamics of 1000 cells were averaged out.

### Nucleation of actin monomers

The process of forming actin trimers, which are essential for treadmill to begin, is called nucleation (Figure 1). Two ATP attached actin molecules (ATP_Actin) molecules collide to form actin dimers (D_1_), which are unstable and quickly self-dissociate into two ATP_Actin monomers (26, 27). However, this binding in presence of formin (FH) results in the formation of stable actin dimers (D_2_). The dimers can quickly be converted into the stable trimers (T_1_ and T_2_) upon immediate availability of additional ATP_Actin molecules in the proximity (26–28). FH consists of two protein domains where one helps attracting profilin attached ATP_Actin (ATP_Actin_Profilin) and the other one provides platform for ATP_Actin molecules to bind with (29, 30). The primary role of FH is considered to speed up the nucleation process by stabilizing dimers and trimers (30). Along with the FH, Capping and Profilin also play an important role in nucleation (30–33). The profilin binds to the ATP_Actin molecules, which are substrate for the dimerization, and forms ATP_Actin_Profilin molecules. This suggests that profilin prevents dimerization by reducing free amount of ATP_Actin molecule in the system (33). However, when the FH is attached to actin dimers (D_2_), profilin actually promotes the formation of trimers (29, 33). Interestingly, the capping proteinis found to compete with FH for bindingwith actin dimers and trimers (34, 35).

**Figure 1.**
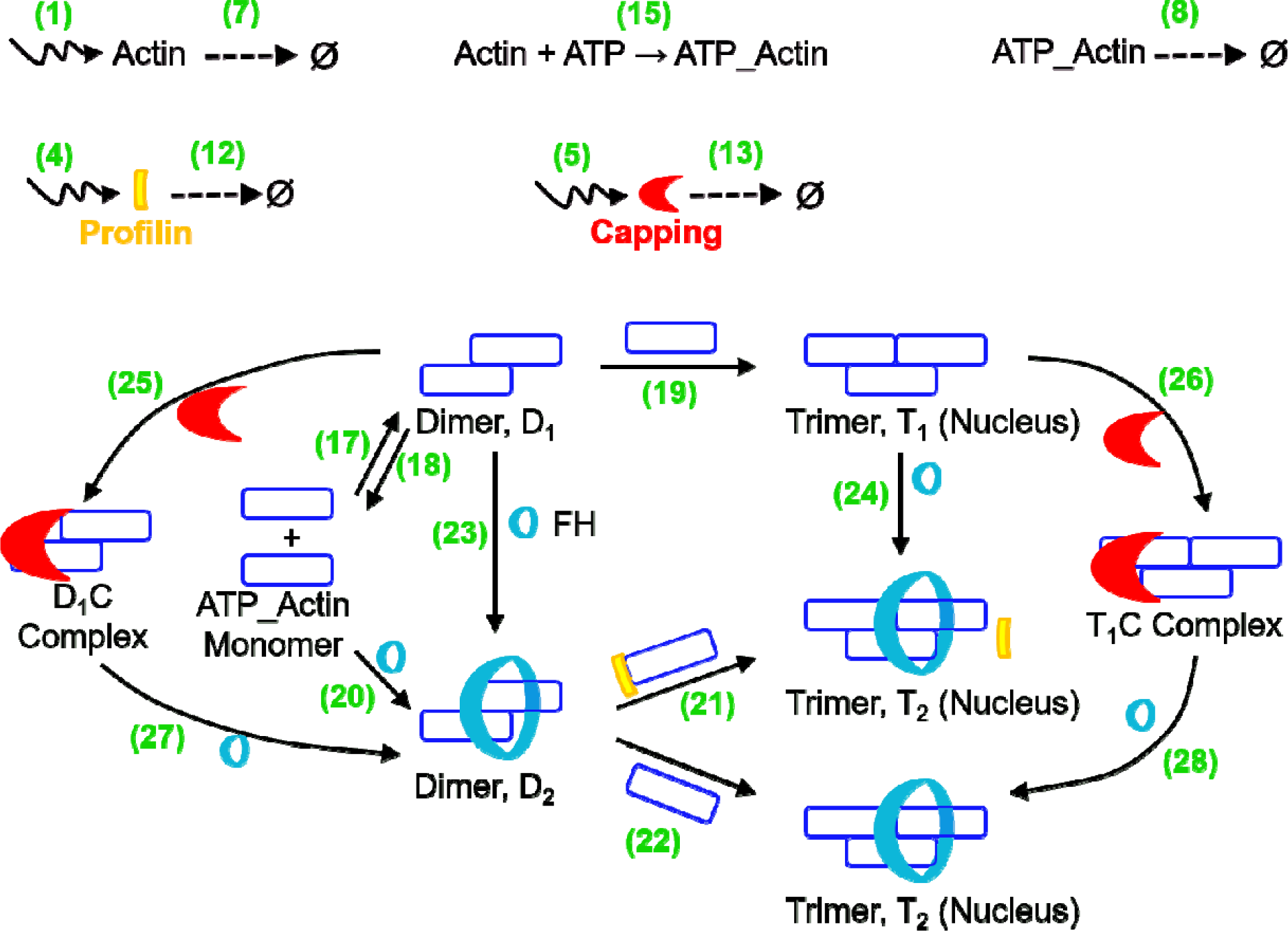
A schematic representation nucleus formation in presence and absence of formin (FH).Each biochemical reaction is marked numerically and number corresponds to the respective ODE mentioned the sub-section “model equations”.

### Elongation of actin filament

The dynamics of actin treadmilling, which leads growth in filament size, depends on the rates of polymerization and depolymerization of actin monomers at the ends of actin filaments. Actin filament elongates when the addition of actin monomers is faster than removal in equilibrium and this process is known as elongation (Figure 2). The process of filament elongation is known to be controlled by many regulators such as profilin, ADF (actin depolymerizing factor), CAP (cyclase associated protein), capping, and coronin(1, 2, 15). Uncommonly, in some species (ex. *Xenopus laevis)*, Arp2/3 is required to form an extensive branched organization of actin filaments in (36). This branching of filaments is not observed in *Plasmodium* (2, 4) thus, such regulators responsible for branching is not included in our model. FH and profilin also play an important role in filament elongation. One FH domain holds filament base and the other one attracts ATP_Actin_Profilin molecules, which results in stabilization of actin filamentsas well as speeding up the polymerization (37).

**Figure 2.**
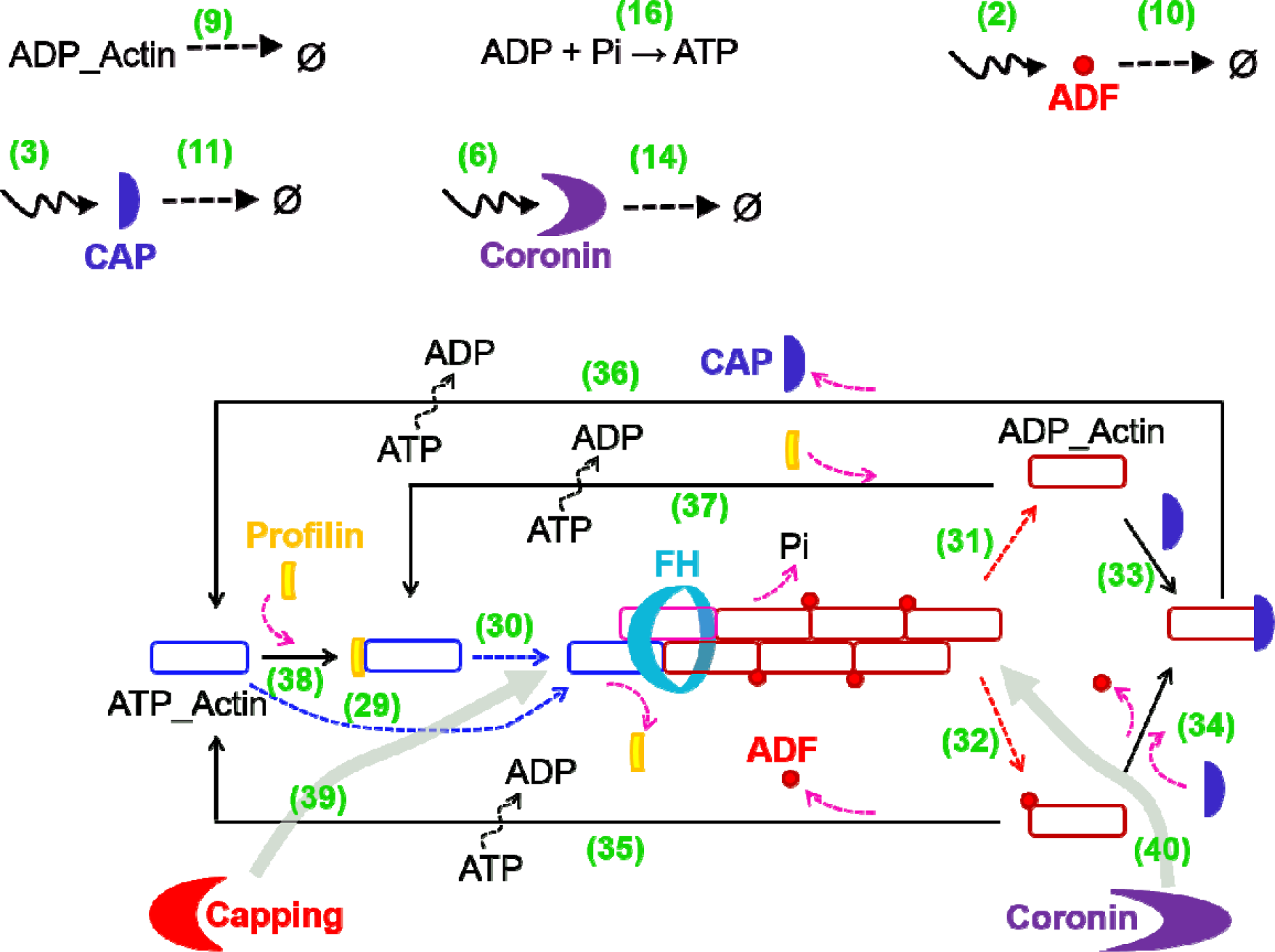
A schematic representation elongation process and balance of molecular species in actin treadmill. Each biochemical reaction is marked numerically and number corresponds to the respective ODE mentioned the sub-section “model equations”.

During the polymerization, the incoming ATP_Actin molecule initiates hydrolysis of ATP attached to the last added actin molecule at the barbed end (Pi still non-covalently attached to ATP) (10, 38) and helps releasing Pi from the penultimate ADP_Pi_Actin molecule (10). This results in the presence of three variants of actin molecules (ATP_Actin, ADP_Pi_Actin, and ADP_Actin) in an actin filament at a time (10). The pointed end of the filament is rich in ADP_Actin molecules, which autodepolymerizes and releases ADP_Actin molecules (10). In addition, ADF/cofilin imposes the strain at the pointed end of the filament and helps depolymerizing actin monomers in the form of ADP_Actin_ADF (10). This suggests that the ADF speeds up depolymerization at pointed end. The ADF with CAP and profilin catalyze nucleotide exchange reactions (39, 40) and convert all forms of ADP_Actin into ATP_Actin. The coronin and capping bind at pointed and barbed ends respectively and abort treadmilling process (41) and dislodge FH from the barbed end(26, 30, 40).

It has been shown *in-vitro* that the actin treadmilling can be performed even in the absence of regulators (26). However, the regulators are thought to be required for the tight and precise control of actin treadmill.

### Filament size, nucleation time, &elongation time

Though the role of each individual regulator in the control of actin treadmill is demonstrated previously (12), the details of biochemical kinetics of treadmilling process and the cross-talk among the regulators are not well understood. In this regard, we perform simulations and explore the dynamics of the treadmill by analysing the filament size, nucleation time, and elongation time in order to understand the kinetic requirements of robust and precise control of treadmill. The filament size is the average length of the actin filaments at steady state, nucleation time is the time required to form actin trimer/nucleus from the actin monomers, and the elongation time is a duration to form full grown filament from an actin trimer. These parameters were analyzed in different variants of treadmill model (as discussed below) to understand the interplay among the regulators.

### Different variants of treadmill model

(i)“WT”, a wild-type variant where all the regulators mentioned above were assumed in the control, (ii)*“No-Regulators”*, a variant, which includes no regulator in the control except capping and coronin, (iii) *“FH (+)”*, where FH is the only regulator in the control, (iv) *“Profilin (+)”*, where profilin is the only regulator in the control, (v) *“FH&Profilin (+)”*, where only FH and profilin present in the control, (vi) *“ADF (-)”*, where all regulators are present other than ADF, (vii) *“CAP(-)”*, where all regulators are present other than CAP, (viii) *“Capping* (-)”,where all regulators are present other than capping.

### Assumptions in themodel

In this model, we only aim to explore a single treadmill in a cell at a time. In order to study the control at single treadmill resolution, only one FH molecule in the cell was assumed thus, can initiate only one treadmill at a time. However, this doesn’t qualitatively change our observation at multiple treadmill resolution. The capping or coronin terminate the running treadmill by dislodging FH from the filament. The dislodged FH is re-used in the next consecutive treadmill. This assumption ensures only one treadmill at a time. Along with the actin molecules, all regulators were assumed to be synthesized at basal rate. All regulators and actin monomers (free, ADP-attached, and ATP-attached) were assumed were assumed to degrade at a constant rate. The actin monomers attached to CAP, profilin, and ADF were not assumed to degrade because of their transient presence and fast inter-conversion.

### Model equations

Basal production of Actin

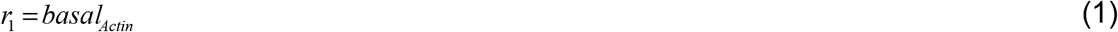

Basal production of ADF

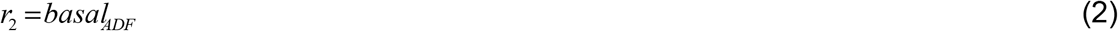

Basal production of CAP

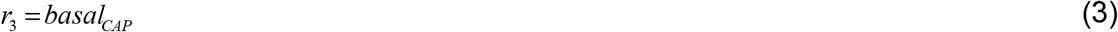

Basal production of Profilin

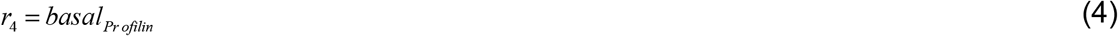

Basal production of Capping

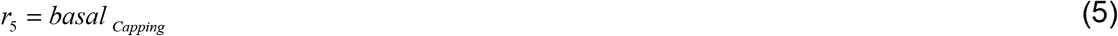

Basal production of Coronin

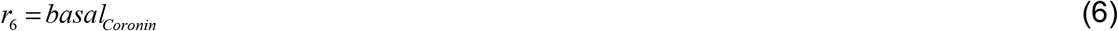

Degradation of Actin

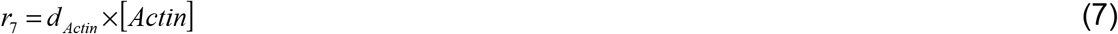

Degradation of ATP_Actin

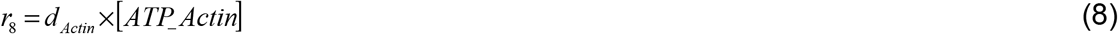

Degradation of ADP_Actin

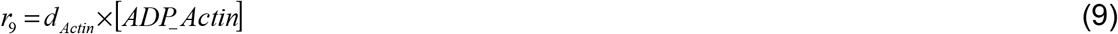

Degradation of ADF

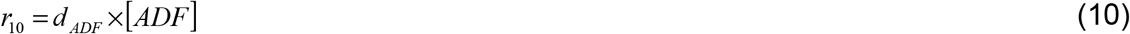

Degradation of CAP

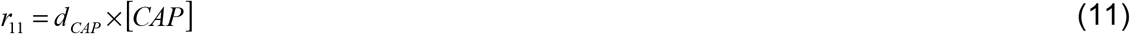

Degradation of Profilin

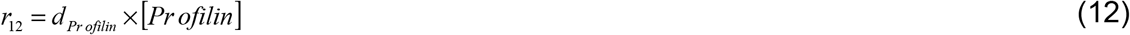

Degradation of Capping

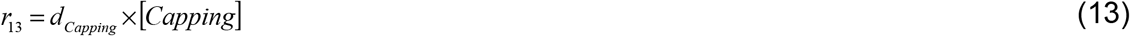

Degradation of Coronin

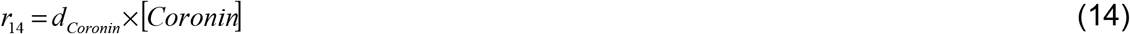

Formation of ATP_Actin

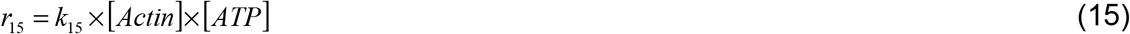

ADP to ATP conversion

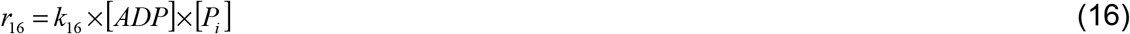

D_1_ formation in absence of FH

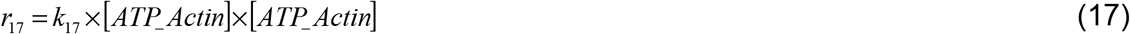

D1 dissociation

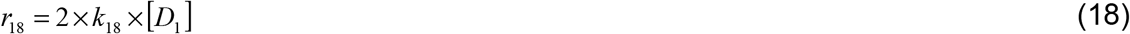

T1 formation from D1

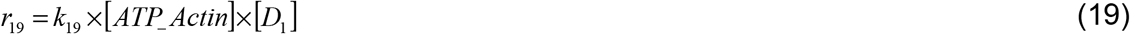

D_2_ formation in presence of FH

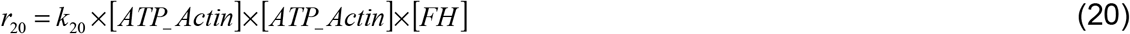

T_2_ formation from D_2_ by ATP_Actin_Profilin

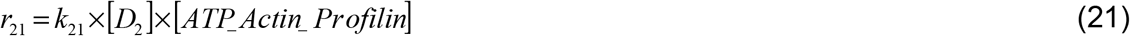

T_2_ formation from D_2_ by ATP_Actin

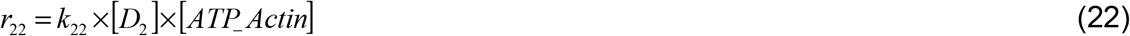

D2 formation from D1 in presence of FH

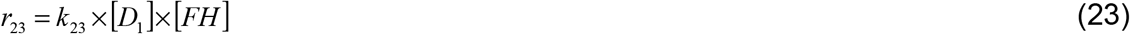

T2 formation from T1 in presence of FH

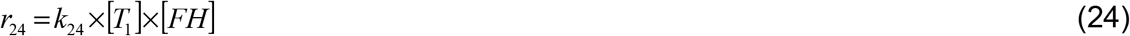

D_1_C formation from D_1_ and Capping

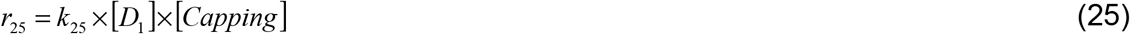

T_1_C formation from T_1_ and Capping

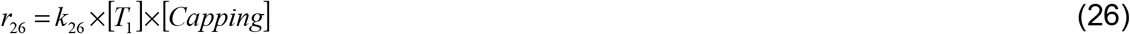

FH dislodges Capping from D_1_C and converts into D_2_

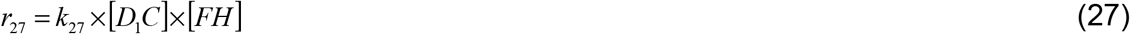

FH dislodges Capping from T_1_C and converts into T_2_

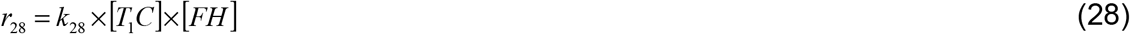

Addition of ATP_Actin at barbed end in absence of FH

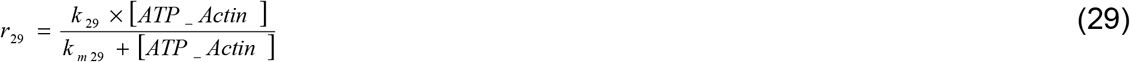

Addition of ATP_Actin_Profilin at barbed end in presence of FH

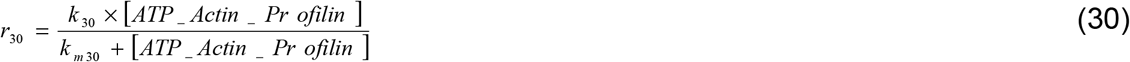

Auto-depolymerization at pointed end

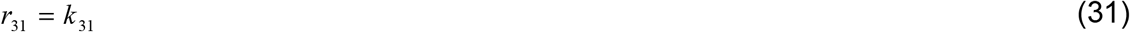

Depolymerization at pointed end in presence of ADF

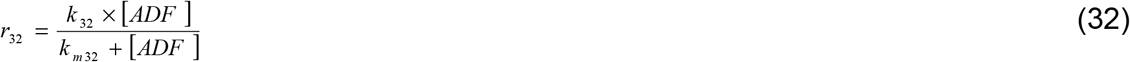

CAP binds ADP_Actin

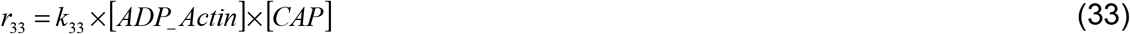

CAP replaces ADF

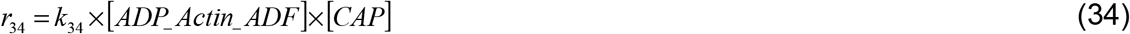

Formation of ATP_Actin from ADP_Actin_ADF

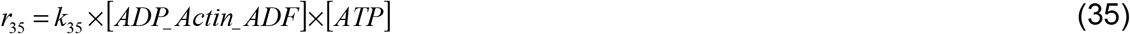

Formation of ATP_Actin from ADP_Actin_CAP

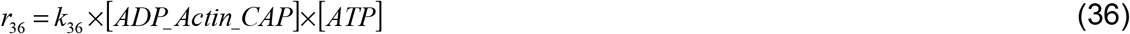

Formation of ATP_Actin_Profilin from ADP_Actin

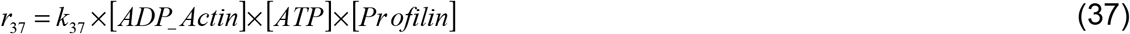

Formation of ATP_Actin_Profilin from ATP_Actin

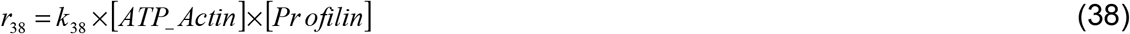

Capping dislodges FH from barbed end

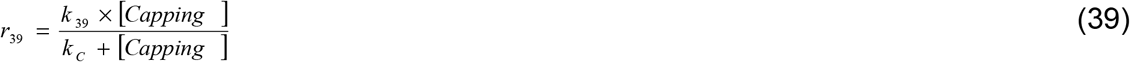

Coronin dislodges FH from barbed end

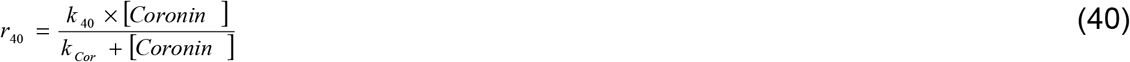

**Table 1.**
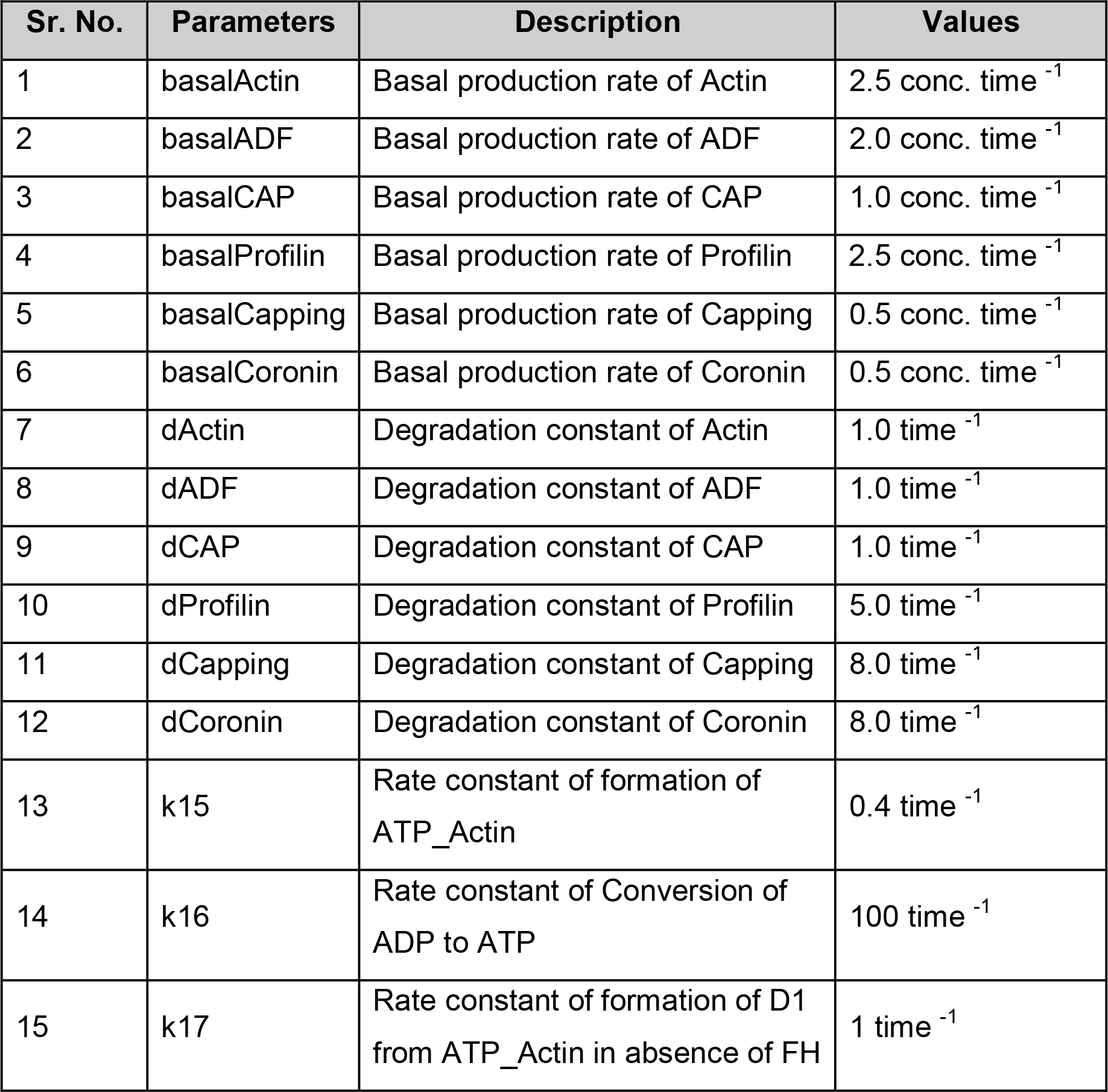
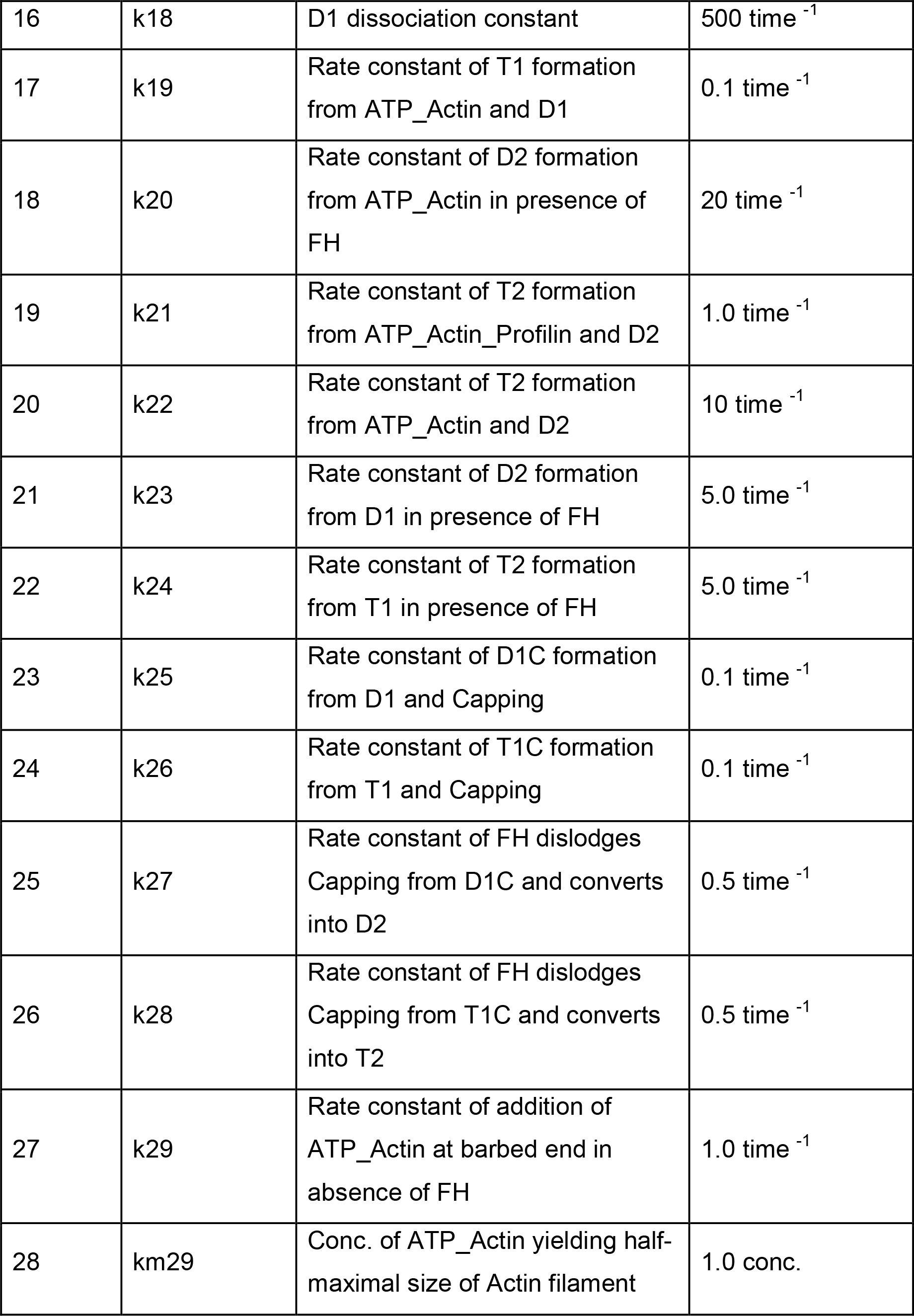
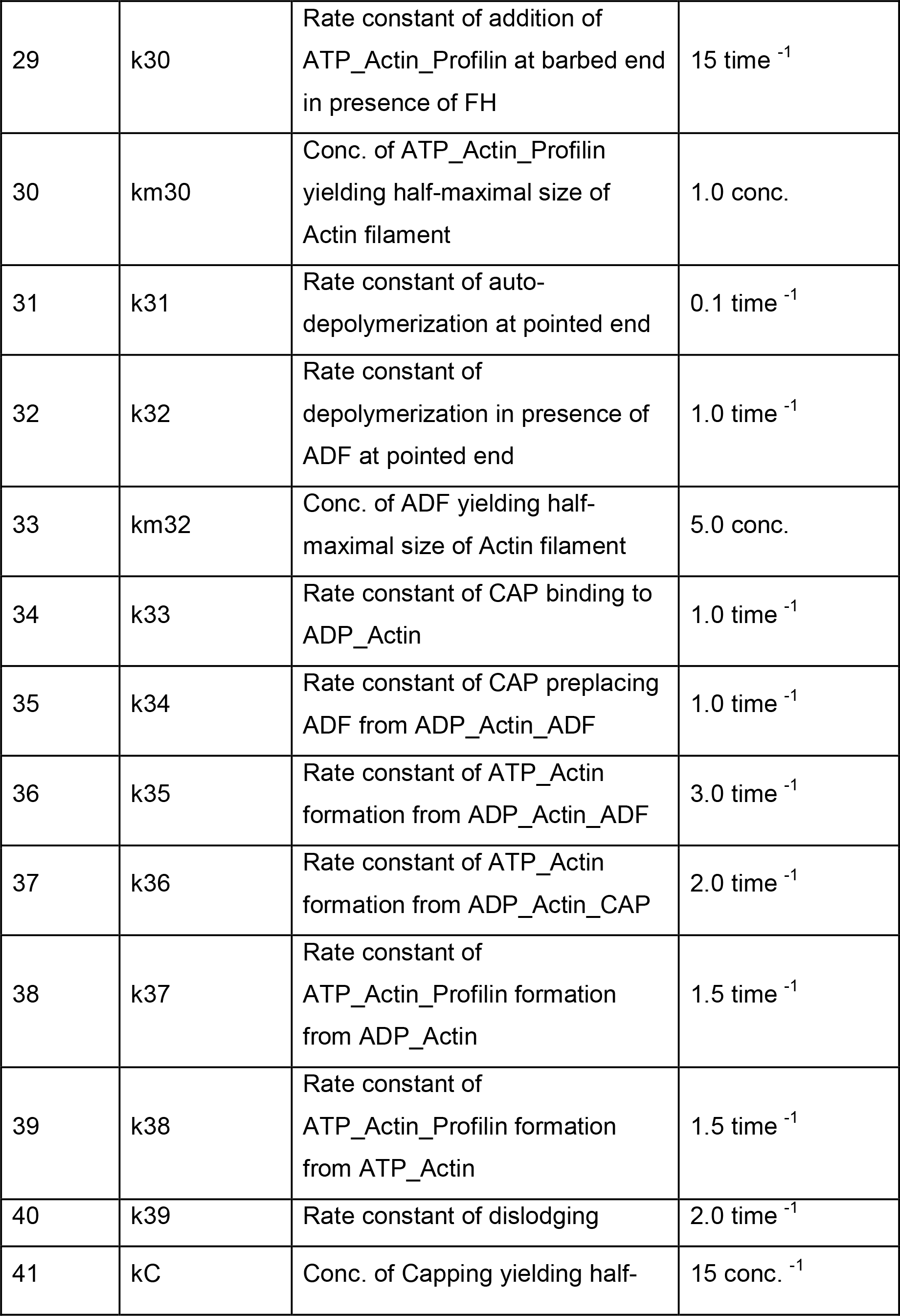
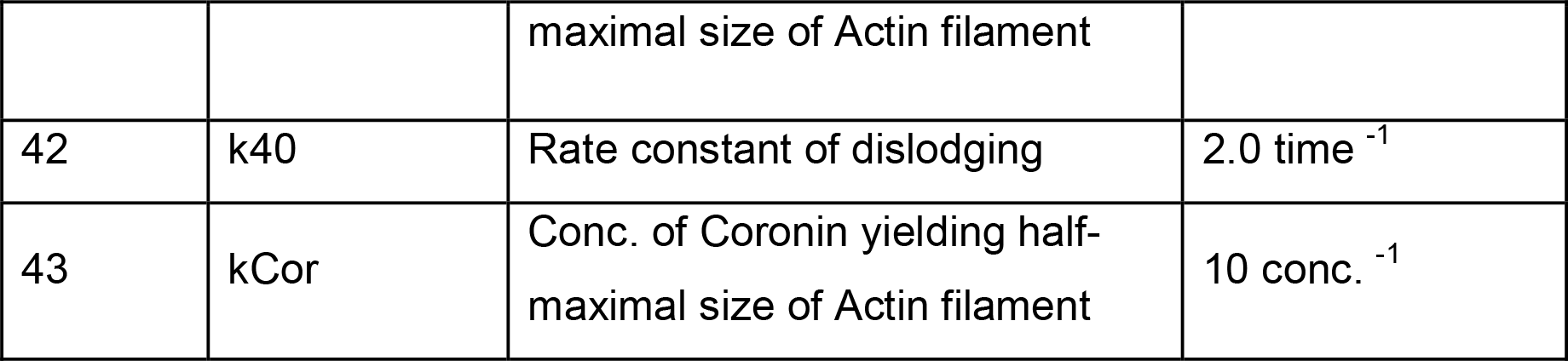
Parameter values used in this study. There is insufficient experimental data to uniquely determine all the parameter values. Therefore, the relative sense of some parameter values has been estimated from previous studies (7, 10, 19–25) and rest are estimated to fit the kinetics.

| Sr. No. | Parameters | Description | Values |
| --- | --- | --- | --- |
| 1 | basalActin | Basal production rate of Actin | 2.5 conc. time<sup>-1</sup> |
| 2 | basalADF | Basal production rate of ADF | 2.0 conc. time<sup>-1</sup> |
| 3 | basalCAP | Basal production rate of CAP | 1.0 conc. time<sup>-1</sup> |
| 4 | basalProfilin | Basal production rate of Profilin | 2.5 conc. time<sup>-1</sup> |
| 5 | basalCapping | Basal production rate of Capping | 0.5 conc. time<sup>-1</sup> |
| 6 | basalCoronin | Basal production rate of Coronin | 0.5 conc. time<sup>-1</sup> |
| 7 | dActin | Degradation constant of Actin | 1.0 time<sup>-1</sup> |
| 8 | dADF | Degradation constant of ADF | 1.0 time<sup>-1</sup> |
| 9 | dCAP | Degradation constant of CAP | 1.0 time<sup>-1</sup> |
| 10 | dProfilin | Degradation constant of Profilin | 5.0 time<sup>-1</sup> |
| 11 | dCapping | Degradation constant of Capping | 8.0 time<sup>-1</sup> |
| 12 | dCoronin | Degradation constant of Coronin | 8.0 time<sup>-1</sup> |
| 13 | k15 | Rate constant of formation of ATP_Actin | 0.4 time<sup>-1</sup> |
| 14 | k16 | Rate constant of Conversion of ADP to ATP | 100 time<sup>-1</sup> |
| 15 | k17 | Rate constant of formation of D1 from ATP_Actin in absence of FH | 1 time<sup>-1</sup> |
| 16 | k18 | D1 dissociation constant | 500 time<sup>-1</sup> |
| 17 | k19 | Rate constant of T1 formation from ATP_Actin and D1 | 0.1 time<sup>-1</sup> |
| 18 | k20 | Rate constant of D2 formation from ATP_Actin in presence of FH | 20 time<sup>-1</sup> |
| 19 | k21 | Rate constant of T2 formation from ATP_Actin_Profilin and D2 | 1.0 time<sup>-1</sup> |
| 20 | k22 | Rate constant of T2 formation from ATP_Actin and D2 | 10 time<sup>-1</sup> |
| 21 | k23 | Rate constant of D2 formation from D1 in presence of FH | 5.0 time<sup>-1</sup> |
| 22 | k24 | Rate constant of T2 formation from T1 in presence of FH | 5.0 time<sup>-1</sup> |
| 23 | k25 | Rate constant of D1C formation from D1 and Capping | 0.1 time<sup>-1</sup> |
| 24 | k26 | Rate constant of T1C formation from T1 and Capping | 0.1 time<sup>-1</sup> |
| 25 | k27 | Rate constant of FH dislodges Capping from D1C and converts into D2 | 0.5 time<sup>-1</sup> |
| 26 | k28 | Rate constant of FH dislodges Capping from T1C and converts into T2 | 0.5 time<sup>-1</sup> |
| 27 | k29 | Rate constant of addition of ATP_Actin at barbed end in absence of FH | 1.0 time<sup>-1</sup> |
| 28 | km29 | Conc. of ATP_Actin yielding half-maximal size of Actin filament | 1.0 conc. |
| 29 | k30 | Rate constant of addition of ATP_Actin_Profilin at barbed end in presence of FH | 15 time<sup>-1</sup> |
| 30 | km30 | Conc. of ATP_Actin_Profilin yielding half-maximal size of Actin filament | 1.0 conc. |
| 31 | k31 | Rate constant of auto-depolymerization at pointed end | 0.1 time<sup>-1</sup> |
| 32 | k32 | Rate constant of depolymerization in presence of ADF at pointed end | 1.0 time<sup>-1</sup> |
| 33 | km32 | Conc. of ADF yielding half-maximal size of Actin filament | 5.0 conc. |
| 34 | k33 | Rate constant of CAP binding to ADP_Actin | 1.0 time<sup>-1</sup> |
| 35 | k34 | Rate constant of CAP preplacing ADF from ADP_Actin_ADF | 1.0 time<sup>-1</sup> |
| 36 | k35 | Rate constant of ATP_Actin formation from ADP_Actin_ADF | 3.0 time<sup>-1</sup> |
| 37 | k36 | Rate constant of ATP_Actin formation from ADP_Actin_CAP | 2.0 time<sup>-1</sup> |
| 38 | k37 | Rate constant of ATP_Actin_Profilin formation from ADP_Actin | 1.5 time<sup>-1</sup> |
| 39 | k38 | Rate constant of ATP_Actin_Profilin formation from ATP_Actin | 1.5 time<sup>-1</sup> |
| 40 | k39 | Rate constant of dislodging | 2.0 time<sup>-1</sup> |
| 41 | kC | Conc. of Capping yielding half- maximal size of Actin filament | 15 conc.<sup>-1</sup> |
| 42 | k40 | Rate constant of dislodging | 2.0 time<sup>-1</sup> |
| 43 | kCor | Conc. of Coronin yielding half-maximal size of Actin filament | 10 conc.<sup>-1</sup> |

## Results and Discussion

### Formin and profilin, are functionally complementary to each other, and together maximize outcome of the actin treadmill

A number of previous studies have demonstrated the individual role of formin (FH) and profilin in the control of actin treadmill (29). The FH is well known to bind with monomeric, dimeric, and trimeric forms of ATP_Actin in order to achieve stability (29). Moreover, in the elongation process, the polyprolene stretch of one of the FH domains specifically attracts the ATP_Actin_Profilin molecules (29). This speeds up the polymerization of actin monomers at barbed end of the filament. Apart from binding with ATP_Actin, profilin also catalyzes nucleotide exchange on actin monomers and converts ADP_Actin into ATP_Actin(42).

FH and profilin are functionally complementary to each other, which means one can’t function without the other’s help. Thus, it would be interesting to see the cross-talk between these two regulators. To do this, we analyzed and compared the wild type (“WT”) treadmill with four different variants *(“No Regulators*”*, “FH (+)*”*, “Profilin (+)*”, and “*FH&Profilin (+)*”) and studied the dynamics of the growth of filament, nucleation time, and elongation time.

In Figure 3A, the variant *“FH (+)*” expressed filaments smaller than that of “WT” and almost equal to that of *“No Regulators*” variant. In absence of profilin, there is no formation of ATP_Actin_Profilin therefore, no additional ATP_Actin_Profilin dependent actin polymerization carried out by FH. Interestingly, no filaments were observed when only profilin was added in the system. Profilin immediately converts ATP_Actin into ATP_Actin_Profilin, which leads to (i) no dimer formation in absence of ATP_Actin and (ii) the access amount of ATP_Actin_Profilin is not in use in absence of FH.

**Figure 3.**
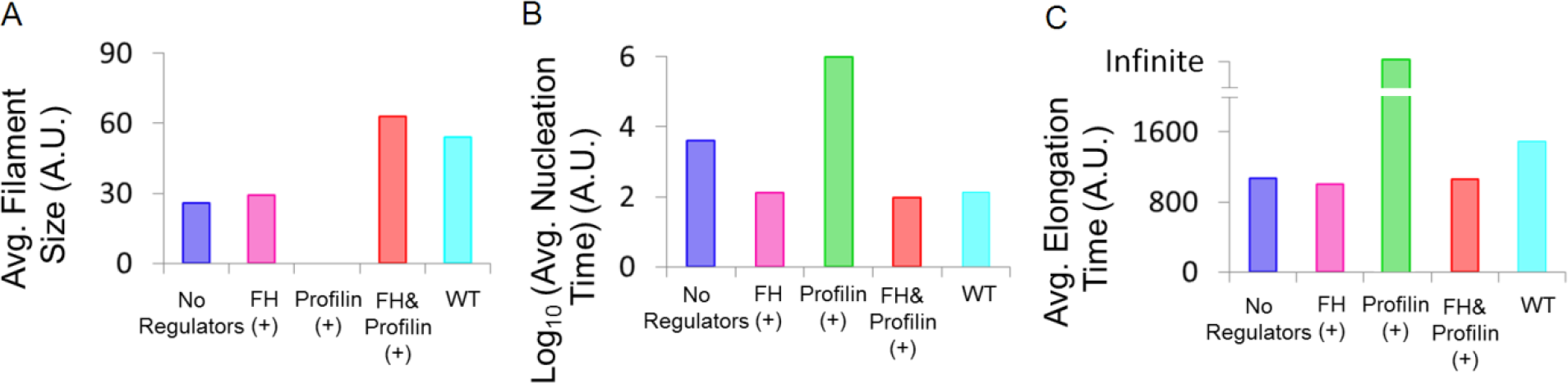
**(A)** Filament size at steady state, **(B)** nucleation time, and **(C)** elongation time were plotted against each treadmill variants.

In Figure 3B, the *“FH (+)*” variant shows reduction in the nucleation time compared to that of *“No regulators*” variant. This is because of the role of FH in stabilizing dimers and trimers of actin, which fastens the nucleation process. The *“Profilin (+)*” variant shows that the only addition of profiling in the system delays dimers and trimers formation by taking away the amount of ATP_Actin.This results in a significant increase in the nucleation time.

Figure 3C clearly shows that the variants *“FH (+)*” and *“No Regulators”* exhibited almost equal elongation time. This is because of nothing can be done by FH solely in the elongation process. However, the variant *“Profilin (+)*” shows infinitely longer time for elongation to complete.

Despite rapid nucleation and elongation in *“FH (+)*” than in “No *Regulators”*, cells were not able to achieve filaments of desired size (exhibited by “WT”). On other hand, *“Profilin (+)*” did not exhibit any filament. However, the variant *“FH&Profilin (+)*” where FH and profilin both together were added in the system was able to optimize the dynamics of filament growth, nucleation, and elongation that was similar to “WT” variant (Figure 3A).

Thus, we predicts a strong interplay between FH and profiling, which synergistically maximizes the outcomes by optimizing treadmill dynamics.

### ADF controls excessive filament growth by delaying nucleation and elongation

The ADF and CAP both catalyze the nucleotide exchange on actin monomers. However, the ADF also plays an important role in depolymerization of actin monomers at pointed end of growing actin filament. How this additional function of ADF can aid control of treadmill? Do ADF and CAP interplay and provide additional feature to the control? In this regard, we analysed and compared *“WT*” with two other variants *“ADF (-)*” and *“CAP (-)*” where ADF and CAP were respectively removed from the “*WT*”.

Upon comparison of “*WT*” and “*ADF (-)*”, it is suggested that ADF plays an important role in the control of excessive growth of filaments by delaying nucleation and elongation (Figure 4). ADF speeds up depolymerization at pointed end and converts ADP_Actin_ADF into ATP_Actin. In absence of ADF, depolymerization of actin monomers was slowed down thus, promoting the size of actin filaments to quickly reach in equilibrium. There is no formation of ADP_Actin_ADF in absence of ADF, which means only ADP_Actin available in the system. ADP_actin molecules can either bind with profilin to form ATP_Actin_Profilin or be converted into ATP_Actin byusing CAP. The presence of both ATP_Actin_Profilin and ATP_Actin at the same time respectively speeds up elongation and nucleation.

**Figure 4.**
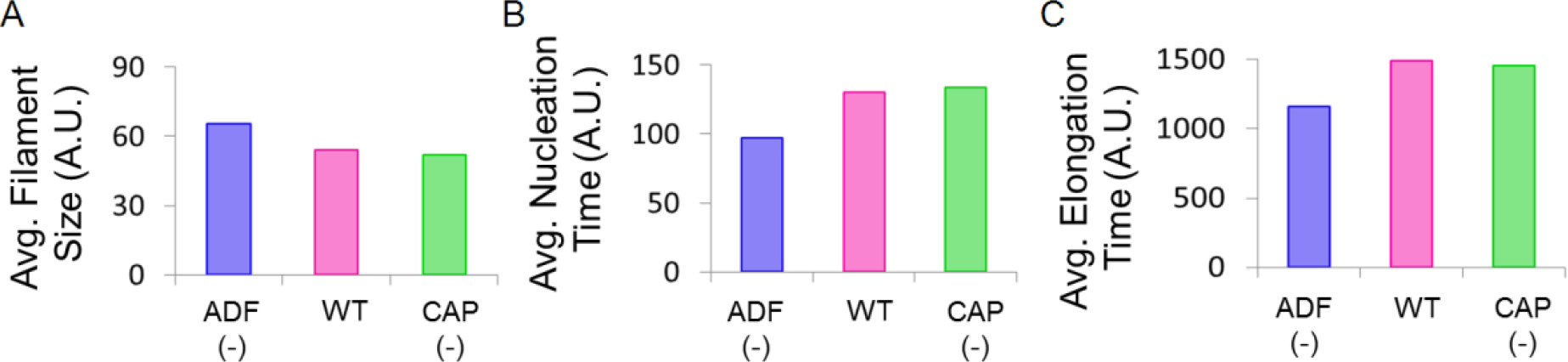
**(A)** Filament size at steady state, **(B)** nucleation time, and **(C)** elongation time were plotted against each treadmill variants.

The comparison of two variants “*WT*” and “*CAP (-)*” (Figure 4) shows no significant effect of CAP in the dynamics of treadmill. The direct role of CAP seems to add an additional way of catalyzing nucleotide exchange. However, it may provide accuracy and robustness to the treadmill by assisting the function of ADF.

### Capping protein enables preserving actin dimers and trimers for next treadmill

Whether capping protein help in terminating or promoting elongation has been debated in a number of reports. This is so because most studies have confirmed its role in termination by binding at barbed end of growing filament (31, 43). During the termination, capping protein also dislodges FH from the barbed end (44, 45). Apart from this, capping protein competes with FH to bind with actin dimers and trimers during in nucleation process. Here, we analysed and compared “*WT*” with *“Capping (-)*” where capping protein was removed from “WT” to understand how capping protein affects the dynamics of actin treadmill.

In the first treadmill performed in absence of capping protein (*“Capping (-)”*), the size of the filaments was increased, the elongation was delayed (Figure 5A&C). Interestingly, in spite knowing the role of capping in nucleation, we did not observe any effect of presence or absence of capping (Figure 5B). To understand this, we further explored three consecutive treadmills (2^nd^ and 3^rd^) in “*WT*” and a variant, which does not allow binding between capping protein and act in dimers/trimers.

**Figure 5.**
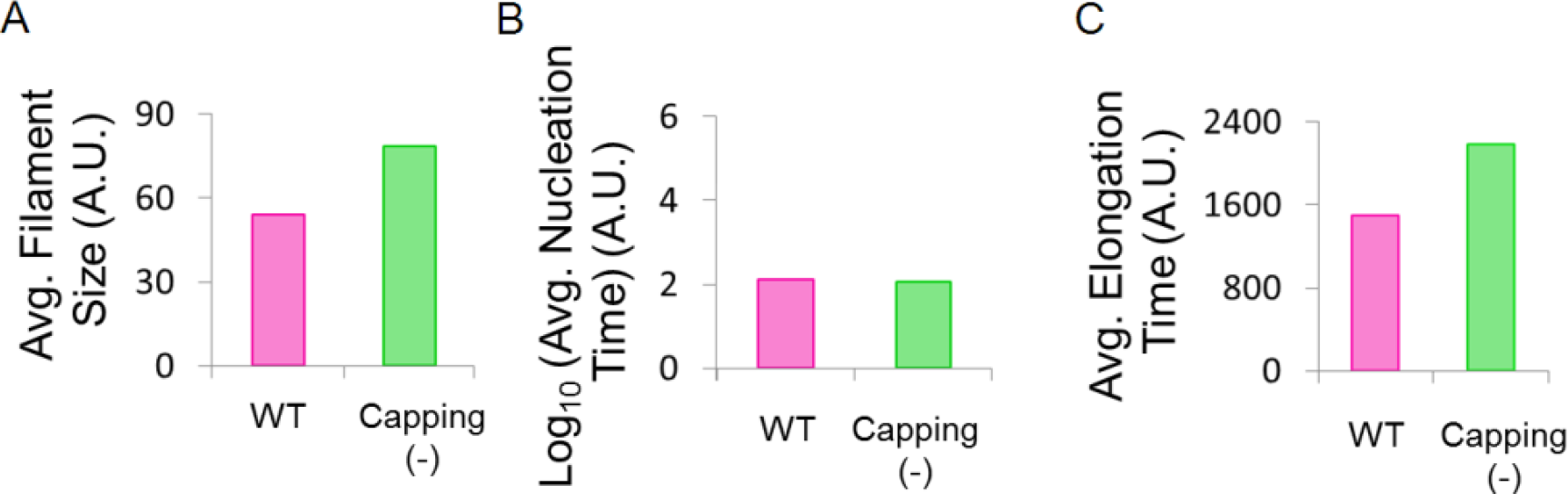
**(A)** Filament size at steady state, **(B)** nucleation time, and **(C)** elongation time were plotted against each treadmill variants.

Figure 6A&C showed no significant difference in filament size and elongation time of the three consecutive treadmills in *“WT*” and the variant. However, the nucleation was constant over consecutive treadmills but significantly differed in the variant (Figure 6B). It appears that the capping protein enables constant nucleation time in the consecutive treadmills without changing filament size and elongation time thus, suggesting a strategy to preserve actin dimers and trimers to initiate actin treadmill at any desired time when the resources are limited.

**Figure 6.**
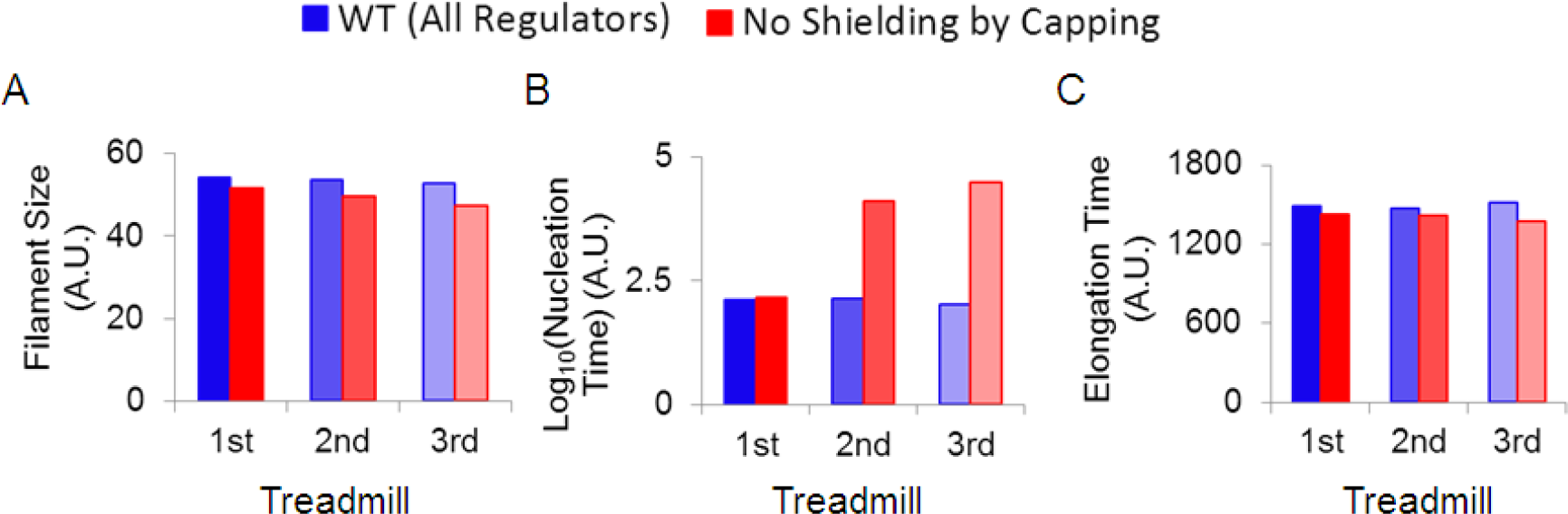
**(A)** Filament size at steady state, **(B)** nucleation time, and **(C)** elongation time were estimated in “WT” and a variant with lost shielding effect capping and plotted against three consecutive treadmills.

### Filament size is maintained below threshold size against fluctuations in actin concentration

By varying the concentration of each regulator, we explored *“WT*” treadmill model and studied the dynamics of actin treadmill control. The degradation constant of each regulator was varied to account the change in the concentration.

As discussed in our previous section, we again observed no effect of CAP protein in dynamics of the filament growth, nucleation time, and elongation time (Figure 7). However, the dynamics of treadmill was observed highly sensitive towards the change in the concentration of capping and coronin proteins. Despite similar functionality, ADF and profilin oppositely affect the dynamics of treadmill.

**Figure 7.**
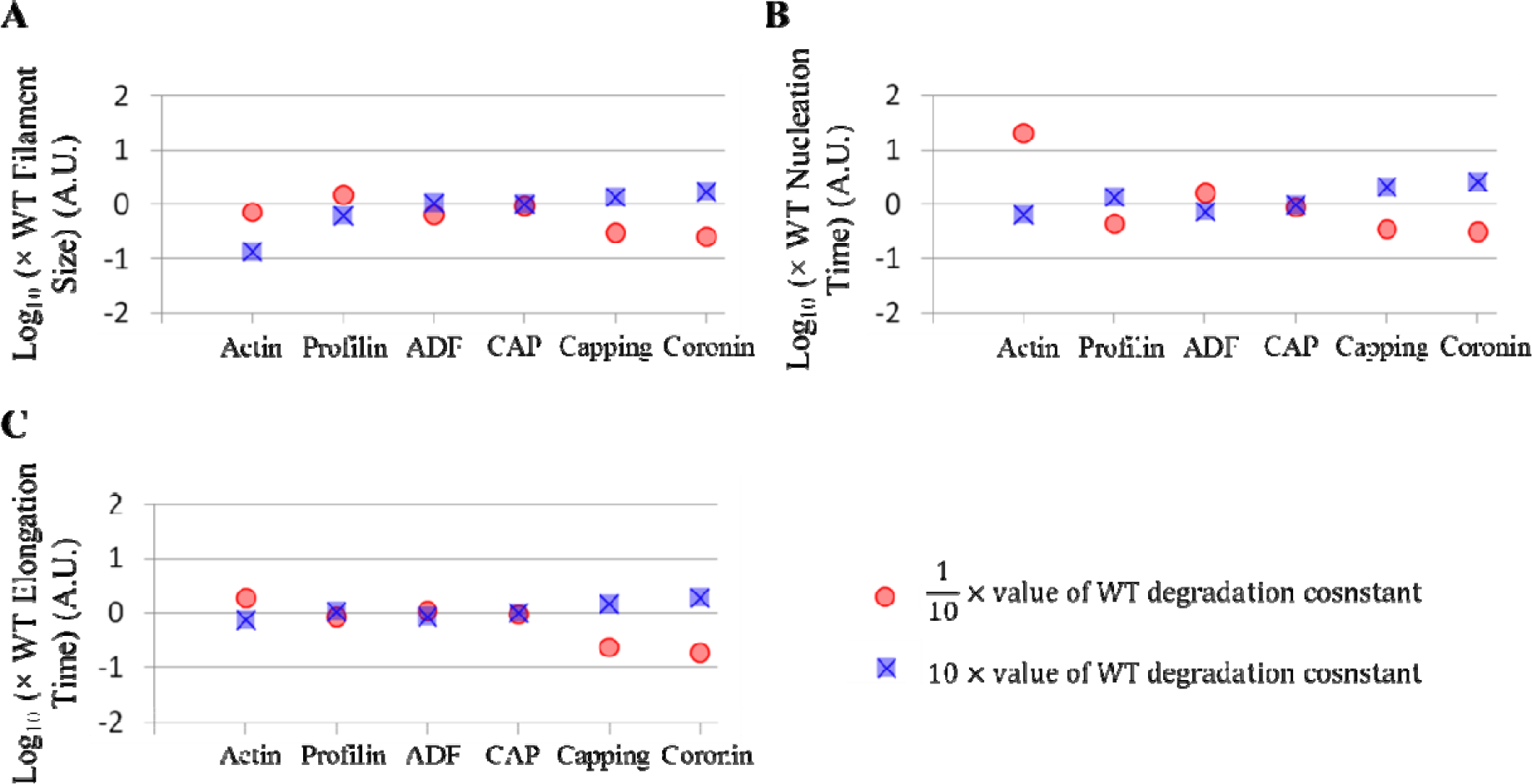
Filament size at steady state, nucleation time, and elongation time are estimated in logarithm against 10 times lower value (red) and 10 times higher of WT degradation constant of each regulator and then plotted.

Surprisingly, the size of the filaments was observed smaller than that of *“WT*” upon both increase and decrease of actin concentration (Figure 7A). This is further explored in the next section.

### Profilin dictates size of the actin filaments in*Apicomplexans*

In agreement with our results (Figure 7), It has been shown previously that the regulation of filament growth should be done in such a way so as to produce actin filaments on and average of the same size in a broad range of variation in actin concentration (10, 11).

Several cellular processes produce and utilize actin monomers and the amount of actin monomers spatiotemporally differed in a cell. This ever-fluctuating amount of actin, which can malfunction the treadmill control such as excessive growth of filaments, must be tackled and strategized to maintain the size of the filaments. Using the *“WT*” model, we deconstructed the treadmill network in a piecewise manner, and elucidated the role of each interaction in dictating constant size of the filaments.

We analysed all the interactions involved in the treadmill (data not shown) and found that the interaction between profilin and ATP_Actin titrated the changes in the concentration of actin and consequently maintain the average size of the filaments (Figure 8, blue curve). Whereas, removing the interaction between profilin and ATP_Actin allowed filaments to grow excessively at higher concentration of actin (Figure 8, red curve).

**Figure 8.**
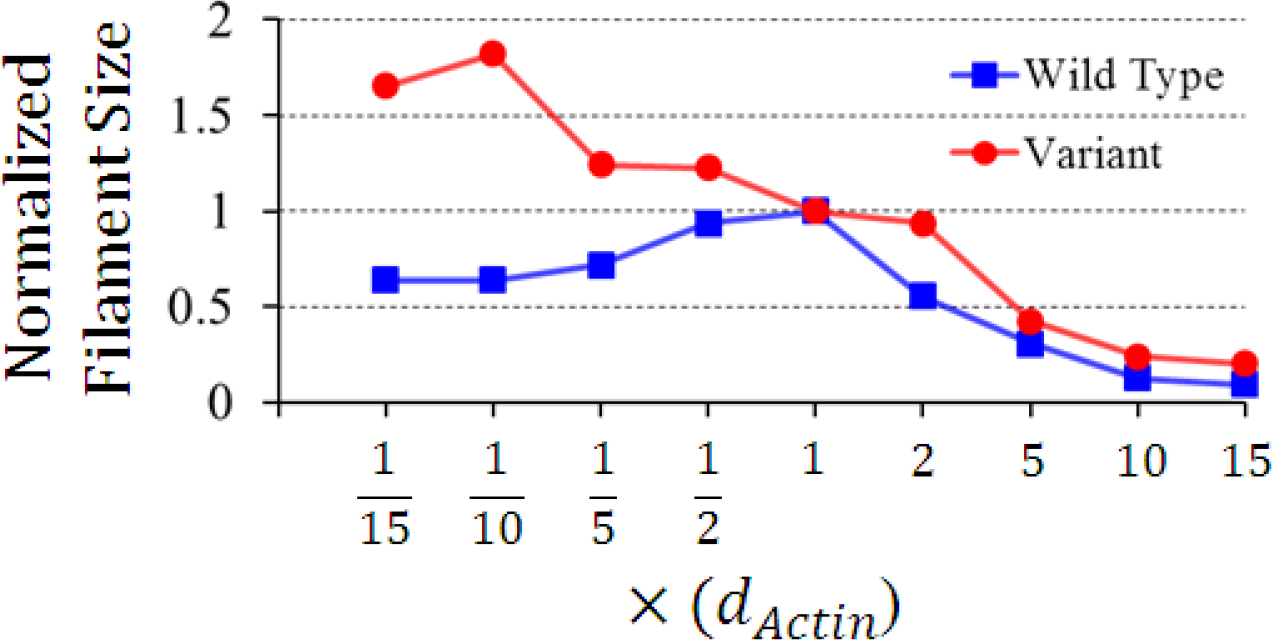
Filament size has been calculated over a wide range of values of actin degradation constant in wild type treadmill (blue) and a variant (red), which does not allow profilin to bind ATP_Actin. X-axis represents filament size at different actin degradation values, which is further normalized by wild type filament size. Y-axis represent a wide range of actin degradation values.

## Conclusions

Reconstruction and a detailed theoretical analysis of actin treadmill control in Apicomplexan helps us understanding its underlying regulatory mechanism and the role of regulators in precise and tight control. The growth of actin filaments can occur even without regulators. However, the regulatory proteins are required for precise and robust control of information processing at treadmill network.

Our results suggest a significant cross-talk between FH and profilin which optimally maximizes the outcome of the treadmill in terms of obtaining the desired filament size by optimizing nucleation and elongation processes. Interestingly, it has been previously reported that CAP and ADF assist nucleotide exchange to convert inactive actin monomers into active form. Though CAP and ADF are known to function similarly, an additional role of ADF in depolymerization of actin monomers at pointed end controls the excessive growth of filaments whereas, CAP only be assumed to achieve accuracy and robustness in treadmill control.

The average size of actin filaments is constant within a species but can vary one to another species (10). *Plasmodium* is such a tiny organism hence; it certainly does not requires large actin filaments to prevent pricking out through its cell membrane (12). How does *Plasmodium* maintain the desired size of the filaments? One of the key observations is that the role of profilin in determining desired size of the filaments in a broad range of fluctuating actin concentration. Another key result is predicting a non-intuitive role of capping protein to preserve actin dimers and trimers for the treadmills so as to ease initiating treadmill when the resources are limited.

While invading the host cells, *Plasmodium* thrives to launch several processes at the same time such as activation of virulence and secretion systems, metabolism, etc. Thus, an excessive cellular energy is required to catalyze these processes at the same time, which can results in a drastic depletion of energy carrier ATPs. In addition, ATPs are also needed to feed in treadmilling process (26). It would be interesting to know that how and what helps *Plasmodium* to recycle ATP?

## Declarations

### Competing interests

The authors declare no competing interests.

### Consent for publication

Not applicable.

### Ethics approval and consent to participate

Not applicable.

### Supplementary files

No supplement attached.

